# A new analytical pipeline reveals metatranscriptomic changes upon high-fat diet in a Down syndrome mouse model

**DOI:** 10.1101/2021.12.16.472765

**Authors:** Ilona E. Grabowicz, Julia Herman-Iżycka, Marta Fructuoso, Mara Dierssen, Bartek Wilczyński

**Affiliations:** Faculty of Mathematics, Informatics and Mechanics, University of Warsaw, Warsaw, 02-097, Poland; Centre for Genomic Regulation (CRG), The Barcelona Institute of Science and Technology, 08003 Barcelona, Spain; University Pompeu Fabra, 08003 Barcelona, Spain; Biomedical Research Networking Center on Rare Diseases (CIBERER), Institute of Health Carlos III, Madrid, Spain

**Keywords:** metatranscriptomics, high-fat diet, Down syndrome, Ts65Dn, microbiome, obesity, contigs

## Abstract

The existing methods designated for metatranscriptomic studies are still rare and being developed. In this paper we present a new analytical pipeline combining contig assembly, gene selection and functional annotation. This pipeline allowed us to reconstruct contigs with very high unique mappability (83%) and select sequences encoding putative bacterial genes reaching also a very high (66%), unique mappability of the NGS sequencing reads. Then, we have applied our pipeline to study faecal metatranscriptome of a Down syndrome (DS) mouse model, the Ts65Dn mice, in order to identify the differentially expressed transcripts. Recent studies have implicated dysbiosis of gut microbiota in several central nervous system (CNS) disorders, including DS. Given that DS individuals have an increased prevalence of obesity, we also studied the effects of a high-fat diet (HFD) on the transcriptomic changes of mice gut microbiomes, as the complex symbiotic relationship between the gut microbiome and its host is strongly influenced by diet and nutrition. Using our new pipeline we found that compared to wild type (WT), Ts65Dn mice showed an elevated expression levels of genes involved in hypoxanthine metabolism, which contributes to oxidative stress, and a down-regulated expression of genes involved in interactions with host epithelial cells and virulence. Microbiomes of mice fed HFD showed significantly higher expression levels of genes involved in membrane lipopolysaccharides / lipids biosynthesis, and decreased expression of osmoprotection and lysine fermentation genes, among others. We also found evidence that mice microbiota is capable of expressing genes encoding for neuromodulators, which may play a role in development of compulsive overeating and obesity. Our results show a DS-specific metatranscriptome profile and show that a high-fat diet affects the metabolism of mice gut microbiome by changing activity of genes involved in lipids, sugars, proteins and amino acids metabolism and cell membranes turnover. Our new analytical pipeline combining contig assembly, gene selection and functional annotation provides new insights into the metatranscriptomic studies.

## INTRODUCTION

Metatranscriptomics is a relatively new field and it poses many challenges, such as identification of species composition from the metatranscriptomic (mRNA) reads as well as identification of active genes. This is a hard task because collections of microorganisms from the gut can exist in different variants, since many bacterial species are evolutionarily closely related or they exchange genes through horizontal gene transfer (Qin et al. 2010). This makes it difficult to definitely claim from which species a gene really comes from (Sankar et al. 2015). Computational methods for differential expression in metatranscriptomic analysis are still not well established and there are multiple approaches found in the literature. Here we applied two known approaches to study the metatranscriptome: mapping to reference genomes and fragment assembly. We compared various mapping methods and tried a handful of *de novo* assembly tools. Then, using the obtained insights, we developed a pipeline to identify, quantify and functionally annotate genes differentially expressed between microbiomes in the different experimental conditions.

In our study, as a proof of concept of our methodology, we analyzed the metatranscriptome from fecal samples from two groups of mice subjected to HFD: Ts65Dn mice, a Down syndrome (DS) mouse model, and their wild-type littermates. Recent studies have implicated dysbiosis of gut microbiota in several central nervous system disorders, including DS (Biagi et al. 2014) and are suggested to be related to cognitive impairment in Chinese population (Ren et al. 2021). DS is the most common genetic cause of intellectual disability due to the trisomy of human chromosome 21 (Lejeune et al 1959; Lott and Dierssen, 2010). DS individuals also present a number of comorbidities, including an increased prevalence of obesity and diabetes (Dierssen et al. 2020). Over the last decade, the gut microbiota has been identified as a potential contributor to the pathophysiology of obesity and related metabolic disorders. The gut microbiota is known to protect gastrointestinal mucosa permeability and to regulate the fermentation and absorption of dietary polysaccharides, perhaps explaining its importance in the regulation of fat accumulation and the resultant obesity.

High-fat diet (HFD) is a common model for studying human metabolic disorders in rodents, as it reflects a trend in modern diets worldwide. HFD leads to body weight gain and usually also to elevated levels of serum glucose, insulin and triglycerides. HFD has been shown to promote binge-eating and compulsive overeating behaviors (Espinosa-Carrasco et al. 2018, De Toma et al. 2018). Many recent studies showed that HFD also promotes changes in the microbiome composition of the subjects (Murphy et al. 2015, Muscogiuri et al. 2019) thought to contribute to obesity through an improved capacity for energy harvest and storage as well as enhanced gut permeability and inflammation. To date, gut microbiome in DS patients has been studied in humans (Biagi et al. 2014; Ren et al. 2021), and in mice (Grabowicz et al. 2020) at the metagenomic, but not the metatranscriptomic level. Here we present a new analytical pipeline combining contig assembly, gene selection and functional annotation revealed DS-specific metatranscriptome profile and show that a high-fat diet affects the metabolism of mice gut microbiome.

## RESULTS

### Transcripts identification

We obtained a total of 83 mln sequencing reads with an average of 10.4 mln reads per sample, in the ribodepleted samples. Using SortMeRNA (Kopylova et al. 2012) we estimated a fraction of ribosomal RNAs ranging from 5 to 32%, with a median of 9% of the reads. Quality of the reads was very good, as assessed with FastQC (Andrews 2010) and therefore it did not require trimming. We used two approaches for identification of the metatranscriptomic transcripts (Fig. 1A). The first aimed to map the sequencing reads on to the genomes of known microbial species and the second to map the reads to the created contigs.

**Fig. 1:**
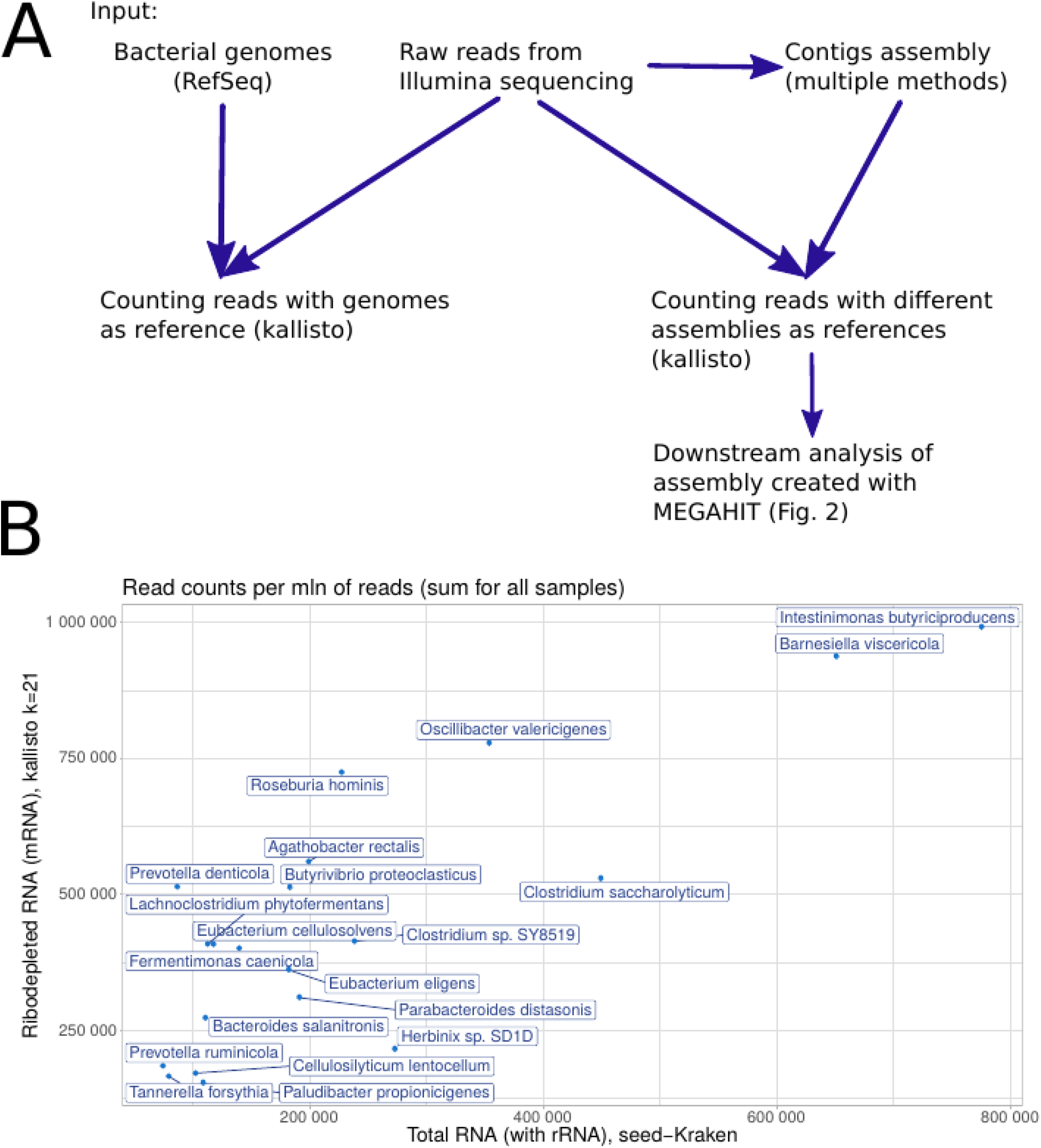
Usage of read quantification tool kallisto for identification of sequences present in metatranscriptomic sample. A) Flow diagram of a pipeline for mapping reads to various reference sequence sets: genomes or contigs. B) Comparison of estimated read counts given by seed-Kraken tool for total RNA samples and Kallisto (k=21) for ribodepleted RNA samples.

#### Mapping rRNA-depleted RNA-seq reads to reference database

As a first step we constructed a database of genome sequences containing 2785 bacterial species with 5242 chromosomal and plasmid sequences downloaded from NCBI/RefSeq (O’Leary et al. 2016) using Clark (Ounit et al. 2015). We treated them as reference “transcripts” and ran Bowtie2 (Langmead and Salzberg, 2012), Minimap2 (Li 2018), fast aligner suited for mapping highly erroneous reads, and Kallisto (Bray et al. 2016), a tool designed for fast quantification of transcripts in complex transcriptomes using RNA-seq data, with different values of parameter k (k-mer size) to find the best setting for further analysis (Table 1). Kallisto (with k=21) gave the highest fraction of mapped reads as it allowed to map uniquely 27% of input reads.

**Table 1:**
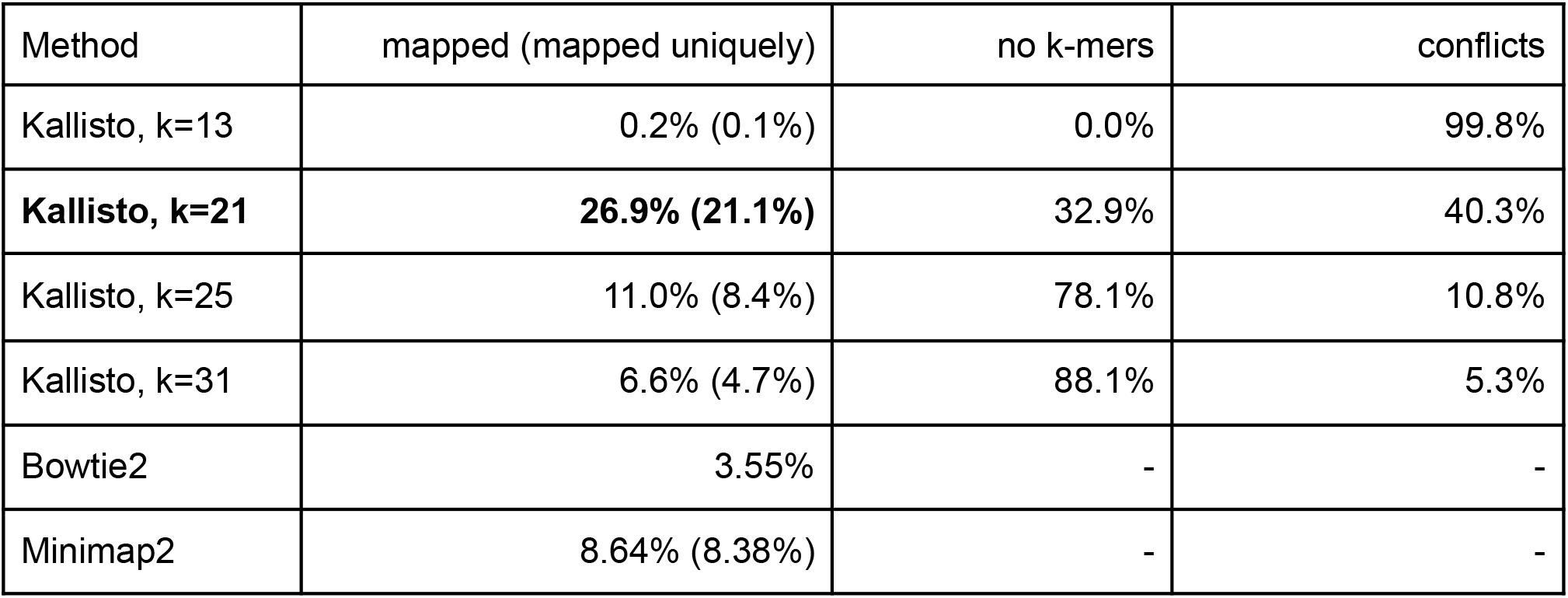
Mean values of percentages of reads mapping to the reference genomes, from all samples. Best results were achieved with Kallisto (with k=21) (bold).

| Method | mapped (mapped uniquely) | no k-mers | conflicts |
| --- | --- | --- | --- |
| Kallisto, k=13 | 0.2% (0.1%) | 0.0% | 99.8% |
| **Kallisto, k=21** | **26.9% (21.1%)** | 32.9% | 40.3% |
| Kallisto, k=25 | 11.0% (8.4%) | 78.1% | 10.8% |
| Kallisto, k=31 | 6.6% (4.7%) | 88.1% | 5.3% |
| Bowtie2 | 3.55% | - | - |
| Minimap2 | 8.64% (8.38%) | - | - |

In order to validate these results, we compared bacterial species abundances assessed with transcriptomic reads (mRNA) with those assessed from total RNA which consists mostly of rRNA (Grabowicz et al. 2020). For assigning the total RNA reads to species we used a seed-Kraken tool (Břinda et al. 2015). Most abundant species according to mRNA RNA-seq results were selected by the number of reads assigned to each species by Kallisto with k = 21. Both methods found the same 20 species to be the most abundant and their abundances were highly correlated (Pearson’s r = 0.81, p-value = 1.5e-05, Fig. 1B).

#### *De novo* assembly of transcripts

In order to achieve a higher fraction of the reads mapping to reference sequences we created contigs, predicted putative genes out of them, and quantified the gene expression levels. Although there are a handful of assembly methods designed for different purposes, we did not find an assembler that was designed for metatranscriptomics. We therefore compared other assembly methods: single-organism de Bruijn graph-based genome assembler Velvet (Zerbino and Birney 2008), which assumes uniform coverage of reconstructed sequences, and Velvet modifications or extensions for transcriptome – Oases (Schulz et al. 2012), which treats repeated regions differently and utilises multiple k-mers –, and for metagenome – Metavelvet (Namiki et al. 2012), which deals with different expected coverages of resulting contigs. We also used string graph method – SGA (Simpson and Durbin 2012), Megahit (Li et al. 2015, Li et al. 2016), a metagenomic assembler using succinct de Bruijn Graphs and IDBA-UD (Peng et al. 2012), a de Bruijn graph assembler, which incorporates multiple techniques to tackle the problem of highly uneven sequencing depth. Contigs were created on the combined reads from all samples with the aforementioned methods. To compare mapping of the reads to the contig assemblies produced with different methods we used Kallisto (k=21) (Table 2). The best result was obtained with Megahit which mapped uniquely over 83% of reads onto over 230,000 contigs. Megahit performed the best also in terms of highest median contig length (median: 543bp). For the subsequent analyses we chose the contigs of length at least 200 generated by Megahit to serve further as a reference for mapping of the reads and quantification of microbial gene expression levels.

**Table 2:**
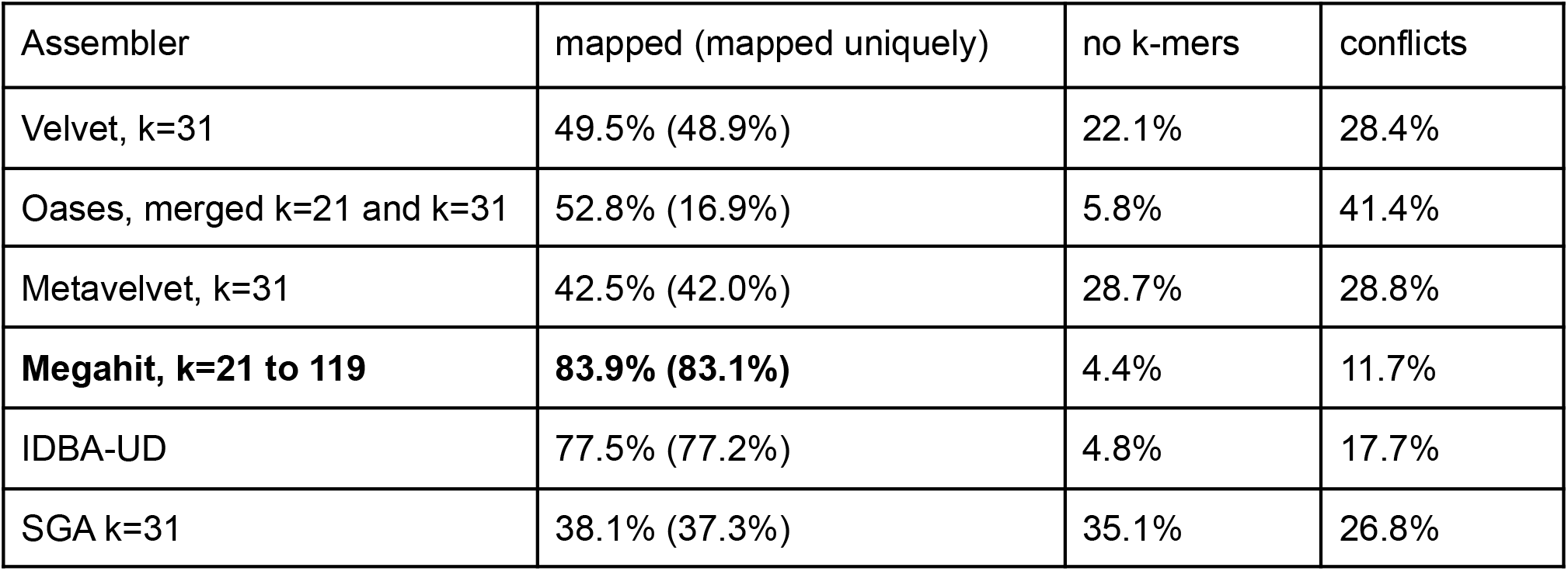
Results of mapping reads with Kallisto k=21 to contigs generated by different *de novo* assemblers. The best result was obtained with Megahit which mapped uniquely over 83% of reads (bold).

#### Counting reads mapping to putative genes in assembled contigs

After creating the contigs from the assembled reads, we developed a pipeline (Fig. 2) which allowed us to identify microbial genes and quantify their expression. From the contigs assembled with Megahit we predicted potential bacterial coding sequences using MetaGeneMark (Zhu et al. 2010). We detected mainly 1 gene per contig (Fig. 3A). We mapped sequencing reads to the sequences of these predicted genes,using Kallisto (k=21) (Bray et al. 2016) obtaining counts for each gene. We reached mean mappability of 67.1% (65.8% uniquely) with 19.9% reads not mapped due to lack of shared k-mer and 13% due to conflicts.

**Fig. 2:**
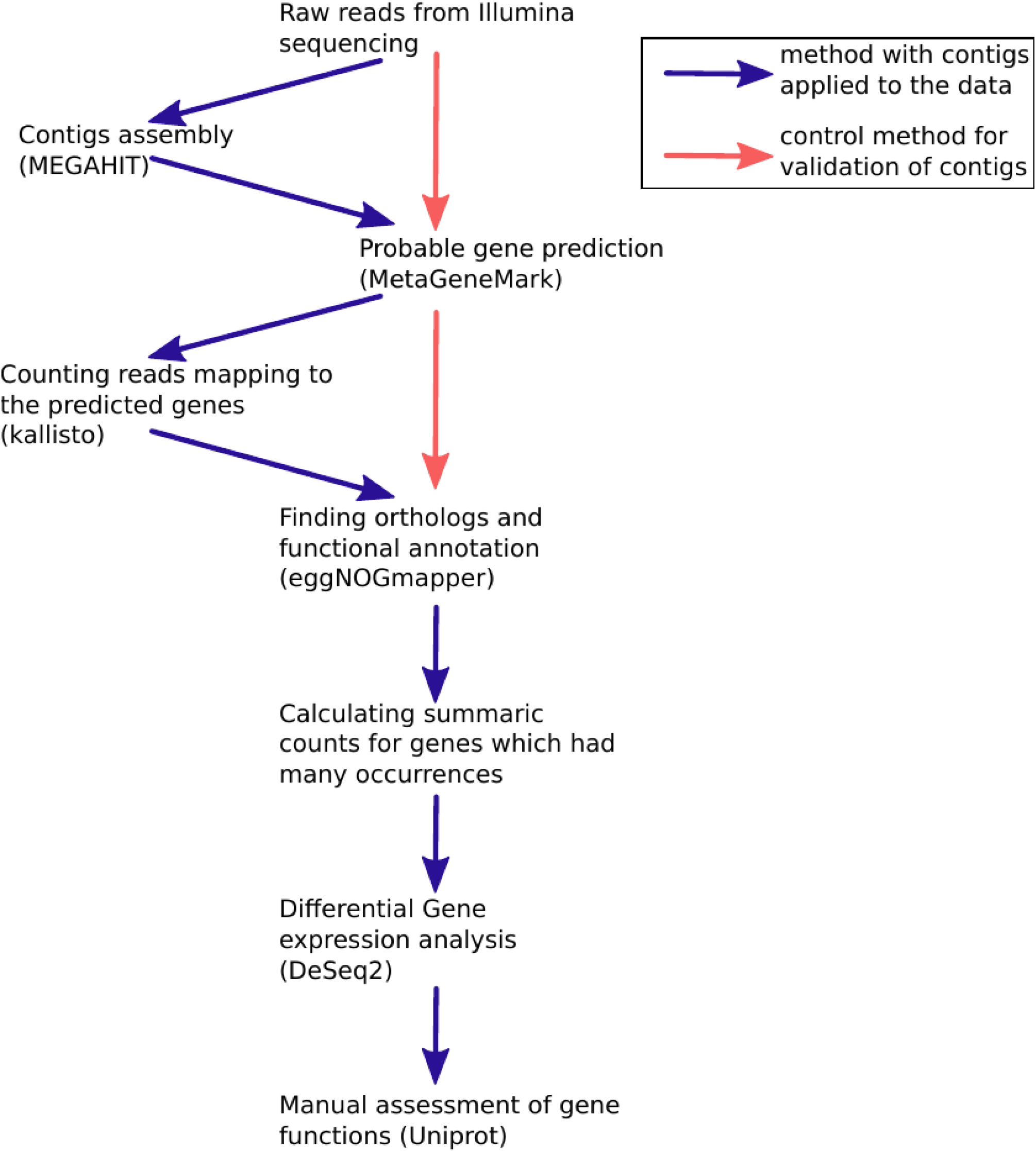
Flow diagram of a pipeline for assessing gene expression. Reads were used to create assemblies using MEGAHIT. From the contig sequences we predicted putative gene sequences using MetaGeneMark and then to these sequences we again mapped using Kallisto (blue path). Alternatively, to control for the correct contigs assembly we predicted putative genes directly from raw reads (pink path). In both paths for the putative genes we found orthologs and assigned the gene names using eggNOGmapper. On the summaric counts of the genes we performed differential gene expression analysis using DeSeq2.

**Fig. 3:**
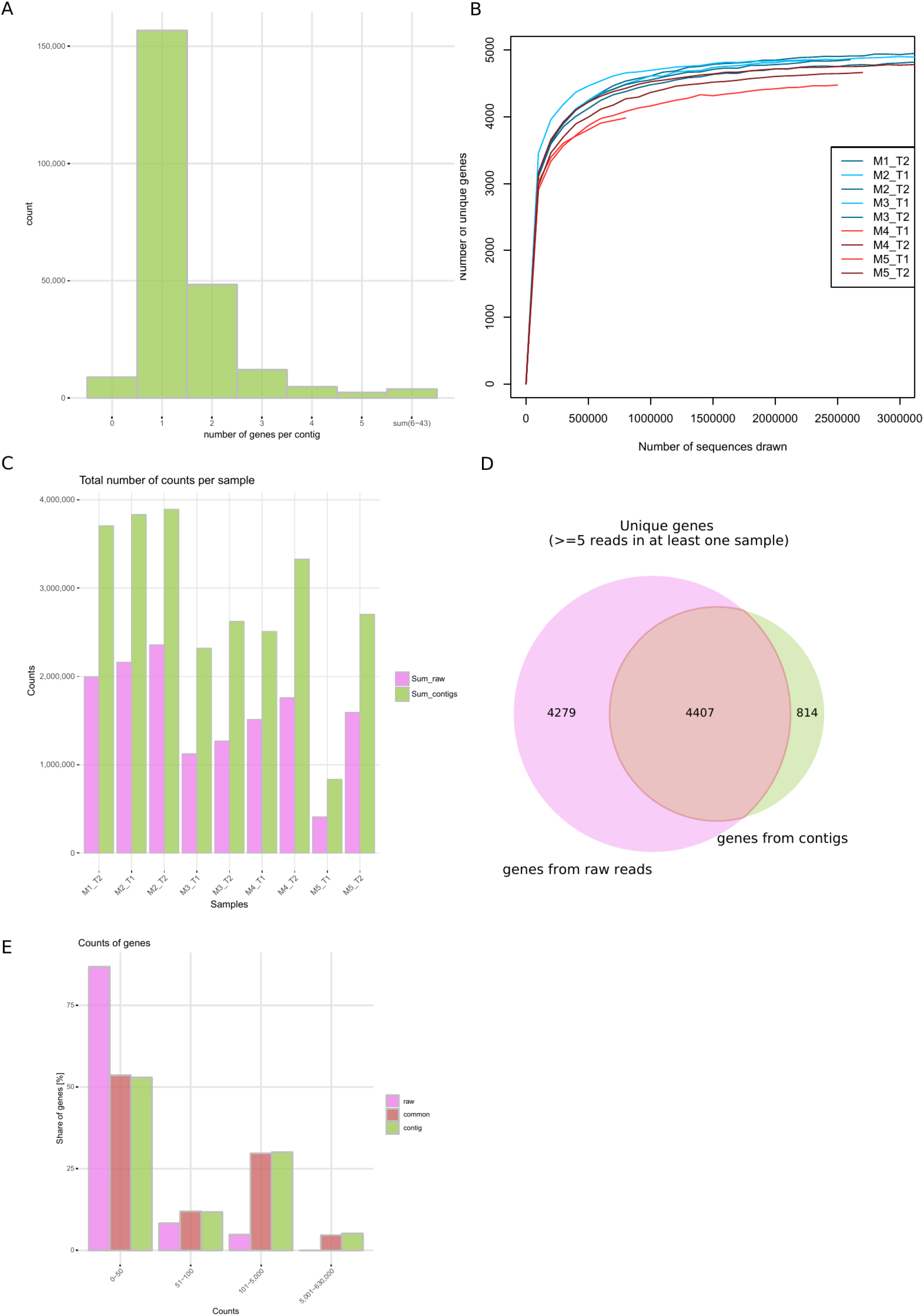
Finding genes in the metatranscriptomic data. A) Histogram of the numbers of predicted genes from contigs. B) Rarefaction curves showing numbers of the predicted unique genes using contigs (the redundant genes counts were summed). In blue: WT mice, in red: Ts65Dn mice; light color: T1, dark color: T2. C) Comparison of numbers of counts mapped to genes for each sample either using contig method or predicting genes directly from raw reads. D) Venn diagram with numbers of unique genes predicted from contigs or directly from raw reads. E) Histogram of percentage of genes to which the low, medium or large numbers of counts were mapped.

Further, we used eggNOGmapper (Huerta-Cepas et al. 2017) to predict orthologs and functionally annotate the genes. On average, we identified the presence of 4827 genes per sample and we found more genes in WT than in Ts65Dn mice (Fig. 3B). For many predicted putative genes, eggNOGmapper assigned the same gene name and function, probably because of the similar gene sequences that were reconstructed from different bacterial species. In order to assess changes of expression of genes among mice of different genotypes or fed HFD, we summed the counts for multiple occurrences of genes with the same names. The sequencing depth used was enough to identify the majority of expressed microbial genes with our pipeline, as the rarefaction curves reached the plateau (Fig. 3B).

To control for the potential misassembly of the Megahit method, we developed an alternative “control” pipeline (Fig. 2, pink-colored path), where the putative gene sequences were predicted directly from raw reads using MetaGeneMark, annotated with eggNOGmapper and the counts for the occurrences of the same orthologs were summed. Of the 20 most abundant genes found by both methods, 12 overlapped between them (Table 3). The method using contigs, however, was able to map nearly twice more reads, therefore making more use of the available data (Fig. 3C). Out of the genes detected both by the method using contigs and raw reads, 46% were the same (Fig. 3D). Importantly, genes identified only by the raw-reads method were mainly low count genes −95% of them had counts of up to 100 reads (Fig. 3E), therefore had a higher probability of artifactual functional assignments. On the other hand, among genes identified by the contig method, 65% had 100 counts or less, 34% had 100 - 50 000 counts and the remaining genes had up to ~630 000 counts (Fig. 3E). Therefore, for the functional analyses we considered only the genes found with the method using contigs.

**Table 3.** Top 20 genes from the rank sorted by the counts found by the method using contigs. (The full list of genes is available in Supplementary Table 1).

| gene | position in rank with contigs | position in rank using raw reads | counts - contigs | counts - raw reads |
| --- | --- | --- | --- | --- |
| FLIC | 1 | 1 | 2392259 | 629853 |
| DNAK | 2 | 2 | 590624 | 618575 |
| GROL | 3 | 3 | 543280 | 428633 |
| HTRA | 4 | 75 | 387582 | 30272 |
| CLPB | 5 | 9 | 351813 | 268179 |
| TUF | 6 | 4 | 326111 | 417798 |
| PCKA | 7 | 5 | 269168 | 339675 |
| HSP | 8 | 42 | 262838 | 48174 |
| MDH | 9 | 14 | 259689 | 149975 |
| PPDK | 10 | 8 | 245001 | 273079 |
| GAPA | 11 | 7 | 244865 | 282700 |
| FLID | 12 | 808 | 240074 | 1709 |
| NIFJ | 13 | 10 | 219253 | 237748 |
| GSIA | 14 | not found | 207437 | - |
| HTPG | 15 | 17 | 204307 | 105897 |
| FRC | 16 | 169 | 203357 | 15405 |
| GAP | 17 | 6 | 202582 | 298039 |
| PG0694 | 18 | not found | 196516 | - |
| OXC | 19 | 697 | 171783 | 2161 |
| SODB | 20 | 60 | 169933 | 36012 |

### Gene expression analysis

#### Characterization of the expressed genes

The gene with most prevalent expression in the microbiome samples was *fliC* (Table 3), which encodes flagellin - the subunit protein which forms the filaments of bacterial flagella which play a role in bacterial motility, attachment, chemotaxis and virulence (He et al. 2012). The rest of the top 10 most abundant genes are involved in processes such as DNA replication (*dnaK, groL, clpB*), protein maturation and degradation of abnormal proteins *(htrA, hsp*), translation (*tuf)*, gluconeogenesis (*pckA*), tricarboxylic acid cycle (*mdh*), and pyruvate metabolism (*ppdK*) (Table 3). Genes from our data were annotated to 24 functional categories of Clusters of Orthologous Groups (COGs). The most abundant one was category ‘S’, which is associated with proteins of unknown function, followed by category ‘R’, proteins with general function prediction only, ‘G’ - Carbohydrate metabolism and transport, and ‘J’ - Translation, ribosomal structure and biogenesis (Fig. 4A).

**Fig. 4:**
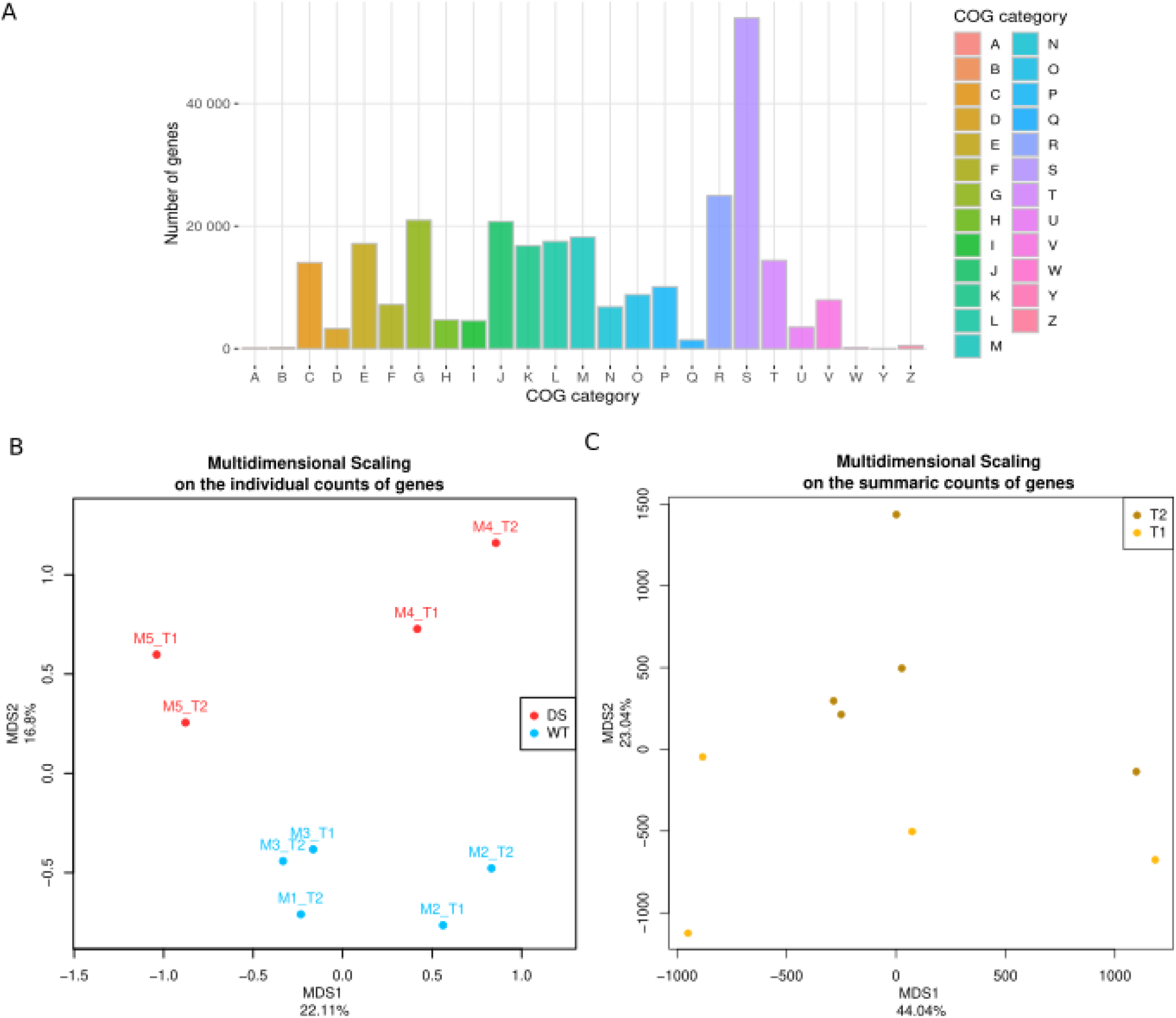
Functional analysis of the genes predicted from the metatranscriptomic data. A) COG categories of the predicted genes. ‘A’ - RNA processing and modification, ‘B’ - Chromatin Structure and dynamics, ‘C’ - Energy production and conversion, ‘D’ - Cell cycle control, cell division, chromosome partitioning, ‘E’ - Amino acid metabolism and transport, ‘F’ - Nucleotide metabolism and transport, ‘G’ - Carbohydrate metabolism and transport, ‘H’ - Coenzyme metabolism, ‘I’ - Lipid metabolism and transport, ‘J’ - Translation, ribosomal structure and biogenesis, ‘K’ - Transcription, ‘L’ - Replication, recombination and repair, ‘M’ - Cell wall/membrane/envelope biogenesis, ‘N’ - Cell motility, ‘O’ - Post-translational modification, protein turnover, chaperone functions, ‘P’ - Inorganic ion transport and metabolism, ‘Q’ - Secondary metabolites biosynthesis, transport, and catabolism, ‘R’ - General function prediction only, ‘S’ - Function unknown, ‘T’ - Signal Transduction mechanisms, ‘U’ - Intracellular trafficking, secretion and vesicular transport, ‘V’ - Defense mechanisms, ‘W’ - Extracellular structures, ‘Y’ - Nuclear structure, ‘Z’ - Cytoskeleton B) Multidimensional Scaling plot based on the individual counts of genes, showing the first two principal components describing variation in the gene expression data between WT and DS model mice (Ts65Dn). The ‘canberra’ distance metric was used. C) Multidimensional Scaling plot based on the summaric counts of genes, showing the first two principal components describing variation in the gene expression data between T1 (day 2) and T2 (day 14) of HFD. The ‘manhattan’ distance metric was used.

Gene expression profiles of our samples showed specificity both for different mice genotypes as well as for the effect of HFD, as the samples segregated into distinct groups in Multidimensional Scaling plots. However, the separation of the samples coming from WT or Ts65Dn mice was visible only when we used the data not pooled for genes (meaning that each gene e.g. *fliC* could have many versions differing slightly from each other, depending on the species it came from) (Fig. 4B), while the separation of samples by the HFD timepoint was visible using the gene-pooled data (Fig. 4C). Moreover, the analysis using non-pooled data revealed the strongest clustering of the samples coming from different time points of the same mouse, indicating that although the microbiome gene expression profiles can change with time and diet they still remain mostly characteristic for each individual. On the other hand, the analysis using pooled data (counts for the same genes, e.g. *fliC* occurring multiple times, were summed) revealed a separation of the samples coming from different timepoints after introduction of HFD (Fig. 4C) indicating that HFD introduced changes in the groups of genes with similar functions, regardless of from which particular bacterial species each gene originated from.

#### High fat diet induces genotype dependent transcriptomic changes

We used DeSeq2 (Love et al, 2014) to identify significantly differentially expressed (DE) genes in microorganisms originating from faeces of wild type and Ts65Dn mice fed HFD. We combined both timepoints (2 days and 14 days of HFD) for this analysis due to the low number of samples. Although we used data from two timepoints our analysis showed a separation between samples from two genotypes in MDS (Fig. 4B). We detected 137 significantly (DeSeq2, p-value<0.05, FDR-corrected) DE genes between the microbiomes of WT and Ts65Dn mice (Fig. 5A). The most prominent categories of DE genes were: transcriptional regulation, membrane transport/turnover, cell wall/capsule biogenesis and carbohydrate metabolism. However, the categories containing DE genes up-regulated in exclusively (or most prevalently) one of the genotypes were: hypoxanthine metabolism being up-regulated in Ts65Dn mice microbiomes, and virulence/interactions with host epithelial cells and transposase activity being upregulated mainly in WT mice microbiomes (Table 4).

**Fig. 5:**
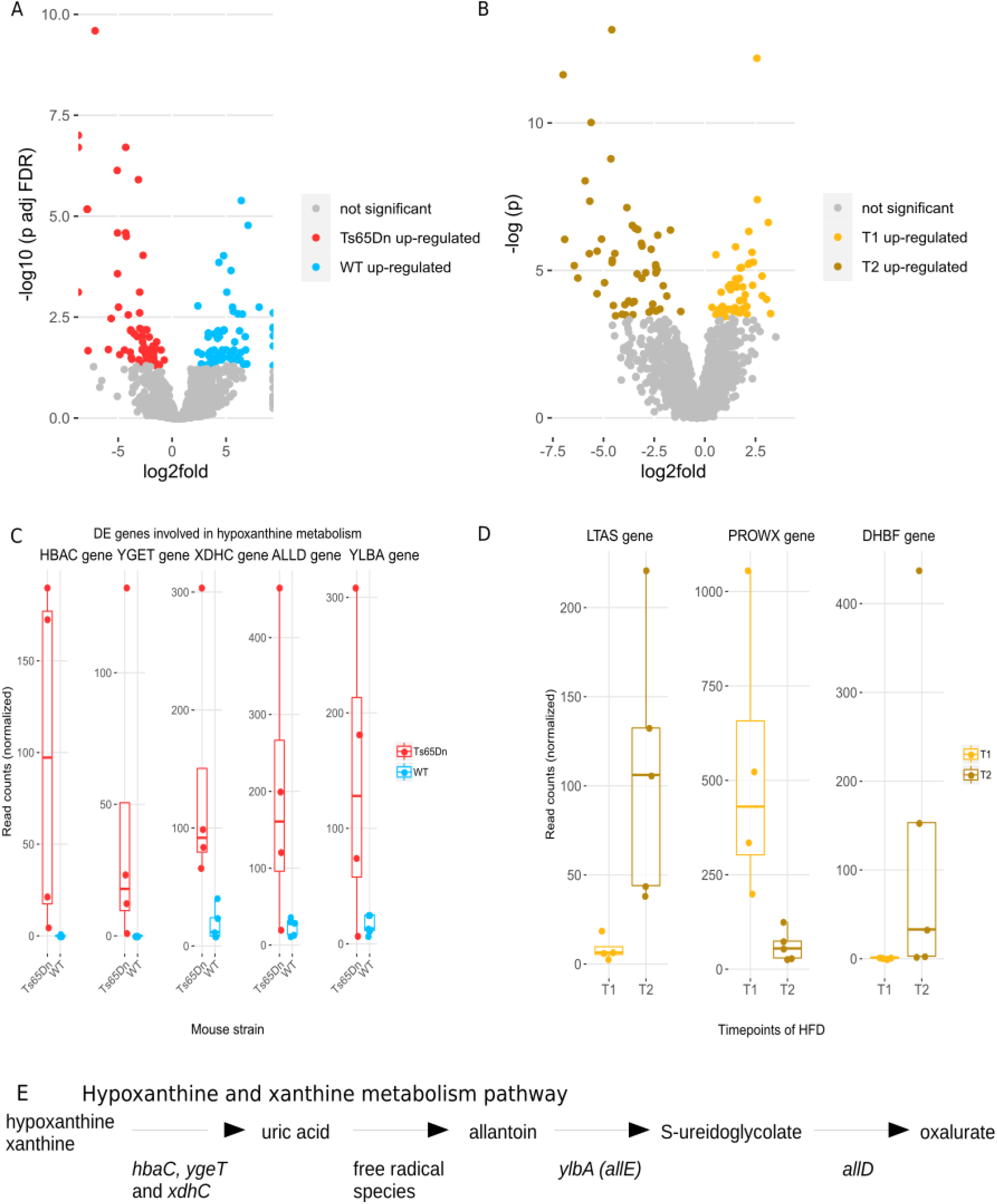
Differential analysis of gene expression. A) Volcano plot representation of DeSeq2 results showing the magnitude (x-axis) and significance (−log 10 of the p-value, FDR corrected; y-axis) for each gene being differentially expressed between WT and Ts65Dn mice. Each dot represents a single gene. Only significantly differentially expressed genes are colored. B) Volcano plot representation of DeSeq2 results showing the magnitude (x-axis) and significance (−log of the p-value; y-axis) for each gene being differentially expressed between faecal microbiomes from T1 (day 2) and T2 (day 14) of HFD. Each dot represents a single gene. Only significantly differentially expressed genes are colored. C) Bar- and dotplots showing expressions of the genes differentially expressed (p-val<0.033, DeSeq2, not FDR-corrected) between WT and Ts65Dn mice, involved in the hypoxanthine/xanthine metabolism pathway. All of them were over-expressed in Ts65Dn mice. D) Bar- and dotplots showing expressions of the 3 FDR-corrected differentially expressed genes between T1 (day 2) and T2 (day 14) of the HFD. E) The hypoxanthine/xanthine metabolic pathway active in the faecal microbiome of Ts65Dn mice.

**Table 4.** Categories of the DE genes (p-value<0.05, FDR-corrected), between wild type and Ts65Dn mice.

|  | WT (up-regulated) | Ts65Dn (up-regulated) |
| --- | --- | --- |
| hypoxanthine metabolism | 0 | *hbaC, ygeT, xdhC, allD, ylbA* |
| virulence genes/interactions with host epithelial cells | *spaA, quiP, nodI, sdrC, phnB, exoT, tetM, VSAL_I0172, enhC* | 0 |
| transposase activity | *CYCMA_1561, DSY0812, THEXY_0761, DACET_0688, RPE_0249, SSQG_02712, BPR_I0156, DALK_0900* | *CPHY_1803, CALNI_0146* |
| transcriptional regulation | *ATP_00026, csm3, treR, LP_2003, LLO_1103, puuR, csiR, sigI, SP_1221, agaR* | *XYLR-1, ocaR_4148, rpiR2, ygeV, gcvR, GYY_09435, yazB, RHOM_13735, nfi, araC8* |
| membrane transport/membrane turnover | *nodI, BL01808, sov, agaF, fecR2, potC, epsK, lktB3, rhaQ* | *xth, BMUL_4061, licB, yhaI, clvE, ptcC, yjbH* |
| cell wall/capsule biogenesis | *spaA, ykvP, VSAL_I0172, murE* | *cap5L, cap5H, wchF, ylbE, mgt, BMUL_2968* |
| carbohydrate metabolism | *cbpJ, bgaA, nanM, rsgI, wcfF* | *pck, bglA, xylBch, amy, aepX, yegV, agp* |

##### Hypoxanthine metabolism

We detected 5 DE genes up-regulated in Ts65Dn mice which are involved in hypoxanthine metabolism (Fig. 5C). Hypoxanthine and xanthine are products of purine metabolism in humans (Žitňanová et al. 2004). They are converted to uric acid and subsequently, in conditions with free radical species present, to allantoin (Grootveld et al. 1987). Three genes involved in the conversion of hypoxanthine to uric acid, namely *hbaC, ygeT* and *xdhC* (Fig. 5C) were up-regulated, and therefore microbiome activity is probably contributing to the aggravation of the oxidative stress in the gut (Fig. 5E). Bacteria are capable of using allantoin as secondary source of nitrogen (Ma et al. 2016) and we could observe that process taking place also in the microbiome of DS model mice: Ts65Dn mice showed elevated levels of gene *ylbA,* also known as *allE,* which converts allantoin to S-ureidoglycolate (Serventi et al. 2010) and *allD,* which provides further conversion to oxalurate (Cusa et al. 1999) (Fig. 5E).

##### Virulence and adhesion to the host

In WT mice we observed 9 up-regulated genes related to bacterial virulence. Two of them *spaA* and *sdrC* help bacteria to adhere to the epithelium of the host. Another gene - *nodI -* is known to be involved in inducing nodulation and symbiosis in plants, perhaps is an ortholog of an unknown gene acting on mammalian gut cells. Two other, *tetM* and *enhC,* provide resistance to antibiotics. *quiP* is involved in quorum sensing (Huang et al. 2006). *exoT* gene is known to be able to change shape of the host cells and help pathogenesis (Sundin et al. 2001). Two remaining genes *phnB* and *VSAL_I0172* encode virulence factors (Uniprot).

#### Transcriptomic changes induced by High-Fat Diet are dynamic

In mice, microbial changes take around 3.5-7 days upon dietary changes to occur and are sustained during weeks (Carmody et al. 2015). To explore the influence along time of HFD on metatranscriptomic profile, we compared the samples collected after the first 48 hours of HFD exposition (T1 = day 2) with the samples collected two weeks later (T2 = day 14) pooling Ts65Dn and WT data. Despite the fact that we used the samples from both genotypes due to the low number of samples the MDS analysis showed a separation between two timepoints (Fig. 4C). There were four genes highly significant DE (p-value<0.05, FDR-corrected) between the T1 and T2 of the HFD: *ltaS, dhbF, proWX* and *OCAR_4032* (Fig. 5D). The most significantly differentially expressed gene was *ltaS,* which was up-regulated in T2 of HFD and encodes lipoteichoic acid synthase (Uniprot). Lipoteichoic acid is a component of the envelope of Gram-positive bacteria and is required for growth and cell division (Gründling and Schneewind, 2007). Another DE gene also up-regulated in T2 was *dhbF*, which encodes dimodular nonribosomal peptide synthetase, which is part of bacillibactin biosynthesis pathway (Uniprot). Bacillibactin is a siderophore - a high affinity metal chelator, which helps an iron-deprived cell to acquire iron and is expressed only when a cell is in low-iron conditions (Lee et al. 2011). HFD has indeed provided twice less iron than standard rodent chow (83 ppm and 159.3 ppm, respectively). The third DE gene was *proWX,* down-regulated in T2, which encodes a glycine betaine/proline ABC transporter (Uniprot). Glycine betaine/proline ABC transporters allow cells to uptake glycine betaine/proline from the environment under high-osmolarity conditions (Csonka 1989). These compounds are used by the cells as osmoprotectants and help survive in stress conditions. The fourth DE gene was an orthologue of OCAR_4032, an uncharacterised gene from *Oligotropha carboxidovorans*. We have also investigated functions of other top 100 most DE genes (although not significantly DE after FDR correction, p-value < 0.033 not FDR-corrected) (Fig. 5B), and found a few categories of functionally related DE genes (Table 5).

**Table 5.** Functional categories often found in the top 100 most DE genes (p-val<0.033), between T1 and T2 of High-Fat Diet.

|  | up-regulated in T1 of HFD | up-regulated in T2 of HFD |
| --- | --- | --- |
| membrane lipopolysaccharides / lipids biosynthesis | *lapB* | *ltaS*, *lcfA*, *lpcA*, *tagA*, *mapA*, *dgkA*, *ugpQ*, |
| virulence | *nanB*, *cpdA*, *cdr* | *iga*, *yhcA*, *mecR*, *SCLAV_0170*, *sle1*, *apf1*, *mapA*, *agrB*, *traF2* |
| transposase | *RV1313C* | *LPG0144, MPOP_0102* |
| protein degradation | *iadA, sspP* | 0 |
| aminoacid metabolism | *hgdB, kamA, kamE, yphF, rbsC-2, kce, guaA, kamD, kdd* | *astA, glxB, tdc, glnN* |
| sugars metabolism | *SP_2141, nanB, porB, sugC, mngB, mtlD, nagE, fucl, rbsK* | *pdhA, pdhB, pdhC, bgaC, lacC, ganA, xfp,* |
| osmoprotection | *proWX, opuCA, sugC* | 0 |
| membrane transport and membrane turnover | *proWX, SCLAV_3941, nirC, SP_1328, oppF3, xth, cdr, yhaH, ygjE* | *cssR, cssS, amt, yhcA, tupA, nagE, opcP, ygiW* |
| transcriptional regulators | *SP_1821, CLS_01740* | *cytR* |
| nonribosomal peptide synthase | 0 | *dhbF* |

##### Osmolar stress genes

Among the top 100 most significantly DE genes, apart from *proWX,* there were two others playing a role in osmoprotection. One of them encodes glycine betaine/carnitine/choline transporter - *opuCA* (Uniprot, Abranches et al. 2006) and the other one was *sugC* which is an ABC transporter complex which is highly specific for uptake of trehalose (Uniprot). Trehalose accumulation in bacteria was found in response to osmotic stress (Csonka 1989). Including *proWX*, all the three genes were down-regulated in T2 of HFD, suggesting that a high-fat diet might cause less osmotic stress on the faecal bacteria.

##### Aminoacid metabolism

Another group of the top 100 most significantly DE genes consisted of 13 genes involved in metabolism of aminoacids: lysine, glutamine, tryptophane, tyrosine and arginine, with most numerous genes metabolising lysine and glutamine (5 genes each). Some of those lead to synthesis while others to degradation of glutamine, while all 5 genes involved in the metabolism of lysine, *kce, kamE, kamA, kamD* and *kdd* catalyze lysine degradation (Kreimeyer et al. 2006) were down-regulated in T2 of HFD. This might indicate that HFD leads to decreased lysine fermentation in bacteria.

##### Genes contributing to bacterial survival

Genes being involved in virulence were mainly up-regulated upon T2 of HFD (9 out of 12 genes). Proteins encoded by these genes help bacteria survive and colonize their host in different modes of action. Some of them provide resistance to antibiotics (*mecR, SCLAV_0170, cdr*), help proper formation of fimbria, which can help in bacterial adhesion to host cells (*yhcA,* Isogai et al., 1988), help assemble pili, which allow genetic code exchange between bacteria (*traF2,* Uniprot) or are involved in quorum sensing system (*agrB,* Uniprot). Other genes such as *cpdA, sle1* or *mapA* alter properties of the cell wall or membrane contributing to the pathogenicity (Uniprot).

##### Lipids metabolism

Despite the existing evidence published by others that gut microbiota are able to express the bile salt hydrolases (BSH) which help them colonize and persist in the gut during gastro intestinal transit (Prete et al. 2020) and that microbial BSH activity changed functions in the gastrointestinal tract, and in the liver of the host (Joyce et al. 2014) we did not observe a significant increase of *bsh* (bile salt hydrolase) (Kaya et al. 2017), nor of genes from the *bai* (bile-acid inducible) operon (Ridlon and Hylemon, 2012) upon HFD. Expression of the *bsh, baiE, baiA, baiB, baiF* and *baiCD* genes was on a very low level (on average <20 reads per sample), and only *baiA* had an average of 600 reads per sample. On the other hand, we found a significant overexpression of genes involved in downstream processing of lipids components such as glycerol-derivatives or fatty acid chains and incorporating them into bacteria’s cell walls, upon prolonged HFD. One of those overexpressed genes - *ltaS* - was most DE between 48h and two weeks exposition to HFD. *ltaS* encodes lipoteichoic acid synthase, which modifies glycerol derivatives and synthesizes lipoteichoic acid (LTA). As shown before (Gründling and Schneewind, 2007) lipoteichoic acid is required for proper cell envelope assembly in *Staphylococcus aureus, Bacillus subtilis* and in other gram-positive bacteria. Other genes overexpressed in T2 of HFD encode enzymes which can modify glycerol derivatives and incorporate them mainly into bacterial cell walls were: *tagA, dgkA* and *ugpQ* (recycling of diacylglycerol, Uniprot). Also, we found overexpression of genes that encode enzymes which can modify fatty acids: *lcfA, lpcA*and *mapA.*

##### Sugar metabolism and membrane transport

Two other categories to which we could assign multiple genes were sugar metabolism and transport across membranes, although we did not see a trend of up-regulation in either T1 or T2 of HFD. Interestingly, three genes from *pdh* operon (pyruvate dehydrogenase), that contains genes encoding multienzyme complexes for the metabolic interconnection between glycolysis and the TCA cycle (Busuioc et al., 2010), were up-regulated in T2. *pdh* operon is predominantly transcribed in the stationary phase, when sugars become a growth-limiting factor. Mice given the choice between HFD and standard chow (SC) systematically preferred HFD. With time, the mice neglected the SC and their meals consisted almost exclusively of HFD. Standard chow is the main source of carbohydrates for mice and after two weeks of HFD they ate less carbohydrates than at the beginning of the diet and therefore activation of the *pdh* operon seems to be a logical consequence.

Interestingly, in T2 of mice fed HFD we observed a significant up-regulation of two components of a two-component regulatory system, named CssR–CssS, found in Bacillus subtilis, which is activated upon so-called secretion stress (Hyyrylainen et al. 2001). The CssR–CssS forms a sensory system controlling misfolding of exported proteins at the membrane–cell wall interface gram-positive bacteria.

## DISCUSSION

Metatranscriptomics is a relatively new scientific area and the tools for analysis of metatranscriptomic data are still being developed and different approaches are being used. The most commonly used approach uses mapping the reads to genomes, either to own in-house databases (Hamilton et al. 2017), or to the publicly available reference genome databases (Reck et al. 2015, Jorth et al. 2014, DeFilippis et al. 2016) or to the genomes of selected microorganisms, which were first identified to be present in the sample (Schirmer et al. 2018). Our results showed, however, that mapping reads directly to genomes yields poor mapping of the reads: only up to 21% of the reads were mapped uniquely (unambiguously). This is probably because of the relatively low coverage of proteins found in the faecal mouse microbiome by the reference databases. Another approach uses *de novo* assembly of contigs and then mapping and counting the reads (Moitinho-Silva et al. 2017, Güllert et al. 2016, Yergeau et al. 2018, Hugenholz et al. 2018, Davids et al. 2016). These authors used a variety of different tools in different combinations, which suggests that there is no consensus in the field about which methods serve best to analyse the metatranscriptomic data. Also, the achieved fractions of mapped reads were sometimes poor - they ranged from 15% to 69% of reads mapped to assembled contigs. In our approach we first assembled contigs out of the reads, then predicted the ORFs and gene coding sequences to which we then mapped back the reads to obtain counts for each gene.

Having tried different alignment tools, we found that k-mer based pseudoaligner Kallisto, a tool already proven to be useful to metagenomic data (Schaeffer et al. 2017), allowed to assign the highest number of reads. Selection of the optimal value of k-mers size was crucial for mappability as it balances between conflicting matchings for small k and no match for large k.

For sequence assembly we also tested a handful of the existing tools. Transcriptomic assembler Oases had low performance because alternative splicing is not an issue with microbial transcriptomics. Metagenomic assembler Metavelvet - a tool designed for assembly of sequences with different coverages - did not improve the mapping rate, because of an unfulfilled assumption about k-mer coverage containing multiple peaks (Suppl. Fig. 1). The reason for a different k-mer distribution is the nature of the metatranscriptomic data, where each species with possibly different abundance produces transcripts at different levels, and those transcripts are short and can be similar between species, thus finding peaks in k-mer distribution is unlikely.

By far, the best results were obtained by another metagenomic assembler, MEGAHIT. To avoid the chimeric contigs we also suggest using some gene identification software such as MetaGeneMark and functional annotation tools like eggNOGmapper downstream of the read mapping to contigs. Thanks to those tools we were able to separate individual genes from larger contigs representing larger parts of bacterial genomes, such as operons or chimeras introduced by assembly errors, pool reads from sequences encoding versions of the same genes and get better predictions of which genes are differentially expressed between samples. Although we found a method to analyse our data, there is still a need for a single comprehensive tool that will combine all steps of analysis: assembly, contig counting, and differential expression analysis. An all-in-one tool could help to use sequenced reads more efficiently, avoiding e.g. “losing” reads that do not map to the produced contigs or found genes and make the process faster.

Our analytical pipeline allowed us to use the majority of the sequencing reads obtained from the microbial metatranscriptomes for identifying differential transcripts. The majority of genes of the effect of a high-fat diet (HFD) on Ts65Dn, a DS mouse model, and WT mice identified in our study, were involved in the basic metabolism: carbohydrate and amino acid metabolism and transport, energy production and conversion, and also in cell growth: replication, transcription, translation and cell wall/membrane/envelope biogenesis. The cell motility category of genes was not the most numerous category of genes, however it contains the gene which was transcribed the most - *fliC*. Unfortunately, despite identification of a large number of transcripts, still a considerable number of predicted genes remained unannotated or poorly annotated.

Congruently with other publications (Woods et al. 2003, Lin et al. 2000) we found that mice fed with a free-choice HFD consumed more energy and gained more weight. According to our previous (Grabowicz et al. 2020) and current results, HFD influences faecal microbiome both on its composition and at the metatranscriptomic level. Among the 100 genes with the largest expression differences, we found genes involved in osmoprotection, which were downregulated after the prolonged feeding with HFD. We can hypothesize that HFD might pose less osmotic stress on the faecal microbiota than standard chow. Such result is in line with a previous publication (Overduin et al. 2014) where the authors investigated the role of substances having different osmotic potential on inhibition of secretion of the appetite-stimulating hormone - ghrelin. They showed that lipids have low osmotic potential, in contrast to glucose which has high.Therefore we conclude that regarding osmotic stress, a high-fat diet might offer the faecal microbiota more lenient growth conditions than a standard chow diet.

Although previous research provided evidence that gut microbiome bacteria are able to express bile salt hydrolases (BSH), which contribute to digestion of fats (Prete et al 2020, Joyce et al. 2014) we found only very low expression of those genes. This might be due to the fact that we studied microbiome of excreted faeces, not the microbiome of the proximal part of the gut, where most of digestion and absorption takes place (Hofmann and Eckman, 2006) and the vast majority of bile acids become actively absorbed by the intestinal epithelium so that less will be available for the microbes downstream. In the faecal microbiome, therefore, we could expect incorporation and metabolism of the earlier-absorbed nutrients as we indeed observed. Upon prolonged HFD, we found a broad group of genes, such as glycerol-derivatives or fatty acid chains, involved in downstream processing of lipids components and in their incorporation into bacteria’s cell walls.

Another effect of HFD we observed was a down-regulation of genes from the *kam* operon, which contains a cluster of genes encoding lysine-fermentation enzymes (Kreimeyer et al. 2007). During the fermentation process, lysine becomes decomposed to acetate, butyrate, and ammonia. Butyrate is known to enhance gut health, by feeding the colonocytes, regulating apoptosis of normal and cancer cells or controlling the intestinal barrier function (Guilloteau et al. 2010). We thus might speculate that HFD diminishes butyrate formation, in line also with other publications, where HFD was associated with lower butyrate production and inflammatory state (Jakobsdottir et al. 2013), while butyrate supplementation in mice fed HFD, prevented obesity (Gao et al. 2009).

Our previous data (Grabowicz et al. 2020), but also others’ (Avena et al. 2009, Sharma et al. 2013) showed that HFD leads to compulsive-like feeding behaviors. There is a growing body of evidence that gut microbiota can produce neurotransmitters such as gamma-aminobutyric acid (GABA), serotonin, norepinephrine, or dopamine (Dinan et al. 2013). In our study, we found >3500 reads per sample of the *gadB* gene encoding glutamate decarboxylase, which produces GABA. The difference in *gadB* expression, however, did not change along HFD consumption, being present in mice microbiomes both at the beginning of HFD and after prolonged HFD. On the other hand, we found a significant increase of expression of the *tdc* gene, encoding tyrosine decarboxylase, which converts tyrosine into tyramine (McGilvery and Cohen, 1948), in mice with prolonged HFD. Tyramine is a neuromodulator, whose receptors are localized within the brain and in peripheral tissues (Xie et al. 2008). Tyramine was found to be involved in sensitization to cocaine in Drosophila, rodents and primates, and facilitates the development of compulsive food intake (McClung and Hirsch, 1999).

Finally, the *dhbF* gene was significantly up-regulated upon prolonged HFD. It encodes a nonribosomal peptide synthase, which is part of bacillibactin biosynthesis pathway (Uniprot) which is expressed only when a cell is in low-iron conditions (Lee et al. 2011). This result reflects well the fact that HFD indeed provided twice less iron than standard rodent chow. Hypoxanthine and xanthine are products of purine metabolism in humans and are converted to uric acid and subsequently, in conditions with free radical species present, to allantoin (Grootveld et al. 1987). In DS patients hypoxanthine and xanthine levels were shown to be lower than in healthy controls while uric acid and allantoin, which are markers of oxidative stress levels, were elevated (Žitňanová et al. 2014). It has been proposed that erythrocyte adenosine deaminase (Becker 1993) or glomerular dysfunction could contribute to hyperuricemia in DS individuals (Nishida et al. 1979). We propose that microbiome activity also could contribute to the oxidative stress as we observed *hbaC, ygeT, xdhC* genes converting hypoxanthine to uric acid being significantly upregulated in Ts65Dn mice, compared to WT mice, considering mice with both short and prolonged HFD feeding. Elevated levels of allantoin comprise a nitrogen source for bacteria being a substrate for their enzymes encoded by the genes from *all* operon (*allD* and *allE*), which we also found up-regulated in Ts65Dn mice, compared to WT mice, considering mice with both short and prolonged HFD feeding.

In conclusion, out of the need to analyse metatranscriptomic data and lack of such tools published we created a new analytical pipeline for metatranscriptomic NGS experiments, consisting of well known tools for contig assembly, gene selection, functional annotation and read quantification. This pipeline was capable of identifying genes changed through prolonged HFD in WT and DS mice. Our results show a DS-specific metatranscriptome profile and show that HFD affects the metabolism of mice faecal microbiome by changing activity of genes involved in lipids, sugars, proteins and amino acids metabolism and cell membranes turnover.

## MATERIALS AND METHODS

### Mice and sample collection

Mice of two strains: WT (n=3) and Ts65Dn (DS model) (n=2) were subjected to a free choice regime of High-Fat Diet and standard rodent chow for 14 days and faecal balls were collected on day 2 (T1) and 14 (T2). High-Fat diet (HF) consisted of commercial pellets of purified diet w/60% energy from fat. HF pellets (58G9; Test Diet ®, USA) contained 22 MJ kg−1 digestible energy (24 % coming from protein, 30 % from carbohydrate and 35 % from fat; carbohydrate is contributed to by 5.4% starch and approximately 6.4 % simple sugars (monosaccharides plus disaccharides). Standard chow (SC) pellets (SDS, UK) contained 10.76 MJ kg−1 digestible energy (17.5 % coming from protein, 75 % from carbohydrate and 7.4% from fat; Table 2). Typically, rodent chow carbohydrate is contributed to by 45% starch and approximately 4% simple sugars (monosaccharides plus disaccharides) as a proportion of total carbohydrates by weight. Faecal samples were collected and directly frozen in −20 C. First, total RNA was extracted from the samples using Total RNA Mini (A&A Biotechnology, Poland) kit according to the manufacturer’s protocol. Then the remaining DNA was digested using 5U of DNAse (A&A Biotechnology, Poland) at 37°C/15 min and then again purified with Total RNA Mini (A&A Biotechnology, Poland). Subsequently, samples were ribodepleted using Ribo-Zero All Bacteria, Illumina protocol. Libraries were prepared using NEBNext Ultra Directional RNA Library Prep Kit for Illumina (New England Biolabs). Validation and quantification of libraries was done using Agilent 2100 Bioanalyzer with Agilent RNA 6000 Pico Kit. Sequencing was performed using 2×100 bp paired-end mode on HiSeq (Illumina) in Genomed SA (Warsaw, Poland) according to the manufacturer’s protocol. 9 samples were subjected to sequencing as from one mouse in the WT group only T1 sample was available. Sequencing reads quality was assessed with FastQC (Andrews, 2010) and the quality of reads from all samples was very high quality, therefore did not require trimming. The fraction of ribosomal reads was assessed with SortMeRNA (Kopylova et al. 2012)

### Bioinformatic and statistical analysis

Codes allowing to reproduce the results shown in this paper and Suppl. Table 1 are deposited at: https://github.com/juliahi/metatranscriptomics_in_DS and https://github.com/juliahi/kallisto

The raw sequencing data used in this study have been deposited in the European Nucleotide Archive (ENA) at EMBL-EBI under accession number PRJEB41352. https://www.ebi.ac.uk/ena/browser/view/PRJEB41352

#### Reference Bacteria sequences

The reference FASTA file was constructed by pasting all bacterial sequences downloaded from RefSeq database on July 25, 2014 using Clark software (Ounit et al. 2015). It contained 5242 chromosome and plasmid sequences from 2785 species.

#### Mapping rRNA-depleted RNA-seq reads to reference database

We ran Bowtie2 (version 2.2.9), in very sensitive mode allowing seed mismatch, and Minimap2 (version 2.16-r922), with default settings for short reads (option -x sr). We also ran our main quantification tool Kallisto (version 0.42.4) with different values of k-mer size parameter k={13, 21, 25, 31}:

Kallisto computes for each read a set of compatible reference sequences by taking some k-mers from the read and computing intersection of sets of reference sequences containing this k-mer (empty sets are omitted). Read is then assigned to one of compatible reference sequences by bootstrap method. In order to find what fraction of reads was mapped uniquely and what was the reason that some reads were not mapped we modified Kallisto to additionally report each reads status:

- size of intersection: if it is == 1 read is mapped uniquely, if it is > 1 mapping is not unique
- size of union of reference sequences: empty union means that read is not mapped because there is no k-mer from read in reference. Otherwise there are conflicting k-mer assignments.

Our modification of Kallisto is available at: https://github.com/juliahi/kallisto

The mappability results in Tables 1 and 2 were presented as means over all samples of the percentages of reads mapping to the reference genomes.

#### *De novo* assembly of reads

Since we did not find an assembler dedicated to metatranscriptomics we compared various other *de novo* assembly methods:

**Velvet** 1.2.10 was run with k-mer size k=31, coverage cutoff cov_cutoff=5, insert length ins_length=200 and expected coverage exp_cov=2

We run **Metavelvet** 1.2.02 on Velvet results for with k=31 and automatic detection of expected_coverage (exp_cov auto):

We run **oases** (version 0.2.09) on two Velvet results, with k=21 and k=31, and read tracking (read_trkg yes) and merged resulting contigs with oases using k=27.

We run **MEGAHIT** (version v1.1.1-2-g02102e1) with k-mer list [21,29,39,59,79,99,119] (default start and step values) and options -m 0.5, -t 12

We run **IDBA-UD** (version 1.1.3) with default settings (i.e. minimum k value = 20, maximum k value = 100, increment of k-mer = 20).

We run **SGA** version 0.10.15 with overlap=31 and k-mer correction. We did not filter out reads with rare k-mers.

In order to find the number of reads mapping to assembled contigs we selected contigs with length at least 200bp (i.e. twice the length of a read) and used them as a reference transcripts set for our modified version of Kallisto, which we run with parameter k=21.

#### Quantification of total-RNA sequences

For the quantification of different species in the total RNA sequences (published in Grabowicz et al. 2020) we used the seed-kraken tool (Břinda et al. 2015).The database for seed-kraken was built based on a collection of sequences from RefSeq (Feb. 2016): Bacteria, Viruses, Fungi, H. sapiens (to exclude possible contaminations) and M. musculus. Sequences needed to have version_status==‘latest’ and assembly_level = “Complete Genome” or assembly_level = “Chromosome”.

#### Assessing gene expression

To assess gene expression from reads mapped to assembled contigs we predicted possible gene sequences from contig sequences produced by Megahit using MetaGeneMark algorithm (Zhu, 2010) - an online tool with default settings. Then we quantified predicted gene sequences using Kallisto (Bray et al. 2016) with k=21. To avoid spurious differential expression calls we removed genes that have an estimated read count of less than 5 in a great majority of samples. In the next step, for the gene sequences predicted from MetaGeneMark we found orthologs using eggNOGmapper v. 1.0.3 (Huerta-Cepas et al. 2017) with the default settings of the online tool in the ‘diamond’ mode. Further with the self-written script we assigned the counts to the orthologues' gene names and for each sample we summed up the counts of genes which had multiple occurrences. Then we performed differential gene expression analysis using the DeSeq2 package (Love et al., 2014) in the R programming language. For visualization purposes (Fig. 5C, D), we normalized the data using fourth quartile normalization.

Additionally, to assess gene expression from raw reads we used MetaGeneMark. We used default settings recommended by the authors of the tool. Then, we also summed up the counts of genes which had multiple occurrences.

#### Differential gene expression

We analysed gene counts with a parametric algorithm from the DESeq2 R package. For T1 vs T2 timepoints we tested only genes with mean of over 20 estimated reads for any of the conditions.

#### Functional analysis of gene expression

For the genes with most reads we searched manually in Uniprot what were their functions. In case of the DE genes comparing T1 with T2 of the HFD we considered 100 genes with lowest p-values. Only 4 of them were significant after FDR correction (corrected p-value<0.05), but all had p-value<0.03 without correction. In the case of DE genes between WT and Ts65Dn mice we considered all 137 genes which had p-value <0.05 after FDR correction. We then manually searched each of the gene names in Uniprot or literature and assigned these genes to functional categories. Some genes were assigned to 2 categories. To some genes we could not assign any function as their annotation was ‘uncharacterized protein’.

#### Results visualisation

Barplots, boxplots and dotplots were generated using R scripting language and libraries: ggplot2 (Villanueva et al. 2016), scales (Wickham and Seidel 2017), dplyr (Wickham et al. 2017a), reshape (Wickham, 2007), tidyr (Wickham and Henry, 2017b), stringr (Wickham, 2017c), egg (Auguie, 2018). Multidimensional Scaling was done using the package ‘vegan’ (Oksanen et al. 2007). Venn diagrams were produced using a python module ‘matplotlib_venn’ (Tretyakov). Rarefaction curves depict numbers of observed species and were generated with a self-written script (R).

## Acknowledgements

The authors thank Pedro Gonzalez Torres and Toni Gabaldon for help with sequencing. IEG, JHI and BW have been supported by the Polish National Center for Research and Development grant [ERA-NET-NEURON/10/2013] and Polish National Science Centre grant [DEC-2015/16/W/NZ2/00314]. The laboratory of M. Dierssen is supported by DIUE de la Generalitat de Catalunya (Grups consolidats 2017SGR926, 2017SGR595). MD also acknowledges the support of Spanish Agencia Estatal de Investigación (PID2019-110755RB-I00/ AEI / 10.13039/501100011033; ERA-NET-NEURON/10/2013 and EMBL partnership), the Centro de Excelencia Severo Ochoa, CERCA Programme/Generalitat de Catalunya, H2020 SC1 Gene overdosage and comorbidities during the early lifetime in Down Syndrome GO-DS21-848077, Fundació La Marató De TV3 (201620-31_MDierssen), EU (JPND HEROES), NIH (Grant Number: 1R01EB 028159-01). MF and MD were supported by Fondation Jérôme Lejeune (2019b – Project #1887) grant.

## Authors contributions

IEG and BW conceived and designed the study. IEG and JHI performed the bioinformatics and statistical analyses, interpreted and visualized results and wrote the original draft; MF and MD organised and performed the behavioral experiments. MF collected and prepared the samples. All authors discussed the results and reviewed and edited the manuscript.

---

**Supplementary Figure 1:**
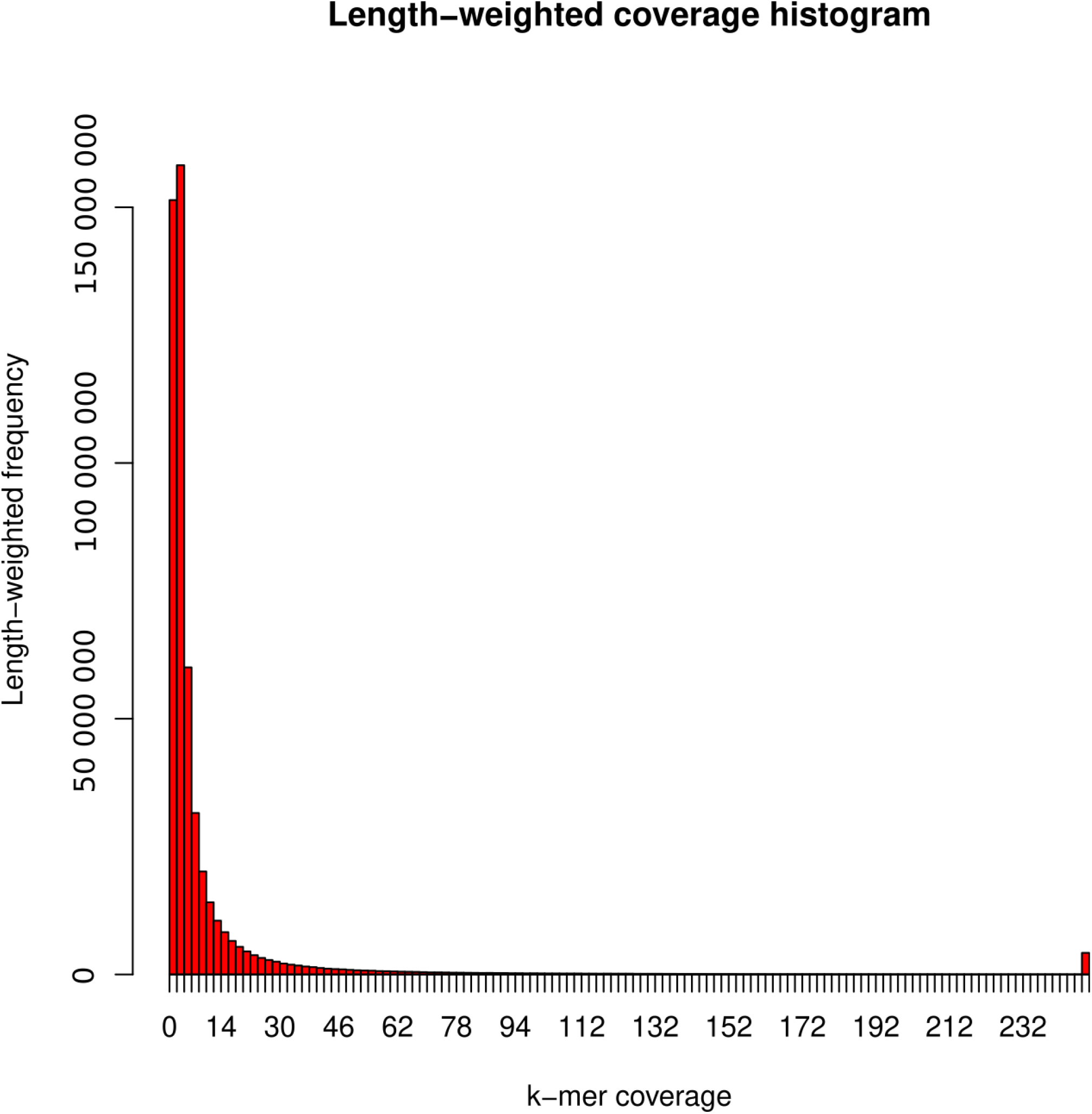
Distribution of length-weight k-mers coverage does not show multiple peaks as expected by Metavelvet to determine coverage peaks.

